# MRI-Trackable Mesenchymal Stem Cells for Oncolytic Therapy: A Clinically Feasible Dual-Labeling Protocol

**DOI:** 10.64898/2026.10.14.759055

**Authors:** Harshan Ravi, Deborah Healey, Neha Nagaraj Damodar, Ellen Ackerstaff, Joy Gumin, Daniel Ledbetter, Frederick F. Lang, Elizabeth J. Shpall, Mayela Mendt Vilchez, Shian-Jy Wang, Christopher Chad Quarles

**Affiliations:** Department of Cancer Systems Imaging, The University of Texas MD Anderson Cancer Center, Houston, Texas, USA; Department of Neurosurgery, The University of Texas MD Anderson Cancer Center, Houston, Texas, USA; Department of Stem Cell Transplantation & Cellular Therapy, The University of Texas MD Anderson Cancer Center, Houston, Texas, USA; MegaPro Biomedical Co. Ltd., Zhubei City, Hsinchu County, Taiwan (R.O.C.); Department of Neuroradiology, The University of Texas MD Anderson Cancer Center, Houston, Texas, USA; Department of Neuro-oncology, The University of Texas MD Anderson Cancer Center, Houston, Texas, USA; Cancer Neuroimaging Research Program, The University of Texas MD Anderson Cancer Center, Houston, Texas, USA; Cancer for Neuroscience of Cancer, The University of Texas MD Anderson Cancer Center, Houston, Texas, USA

## Abstract

Non-invasive imaging of oncolytic virus-loaded mesenchymal stem cells (MSCs) could help evaluate cell trafficking and delivery in cancer therapy. Labeling MSCs with iron oxide particles (IOPs) enables *in vivo* tracking using magnetic resonance imaging (MRI). This study evaluated a dual-labeling strategy that loads MSCs with MegaPro IOPs and the oncolytic virus Delta-24-RGD (DNX-2401; tasadenoturev) and assessed cellular function and MRI detectability. MSCs were labeled with IOPs alone or in combination with the oncolytic virus. We compared cell viability and intracellular iron content among IOP-labeled, dual-labeled, and naïve MSCs, and evaluated differentiation, invasion, and immunophenotype in naïve and IOP-labeled MSCs. We evaluated phantoms containing varying numbers of IOP-labeled MSCs (9 × 10³–1.5 × 10⁵ cells) using R_2_-, R_2_*-, and R_1_-weighted MRI, and generated corresponding relaxation rate maps to assess concentration dependent contrast and detection sensitivity. Iron content was comparable between IOP-labeled and dual-labeled MSCs (9.2 ± 2.6 and 11.6 ± 1.4 pg/cell, respectively), with >91% viability across all groups. IOP-labeled MSCs retained invasion and differentiation potential. At the highest labeled-cell concentration, R_2_*, R_2_, and R_1_ increased approximately 4, 2.5, and 1.5 fold, respectively, relative to the agarose control. The estimated detection limits were approximately 2 × 10³ cells for R_2_* and 6 × 10³ cells for R_2_ and R_1_. These findings demonstrate that dual labeling of MSCs with IOPs and oncolytic virus preserves key cellular characteristics while providing MRI-detectable contrast, supporting further investigation of this approach for image-guided tracking of cell-based cancer therapies.

## Introduction

Delta-24-RGD (DNX-2401; Tasadenoturev) is a promising oncolytic immunotherapy that leverages targeted viral oncolysis and immune activation to treat patients with glioblastoma (GBM).^1–3^ For neuro-oncology applications, oncolytic viruses can be delivered via mesenchymal stem cells (MSCs) as carriers due to their tumor-homing capacity and ability to deliver therapeutic payloads, such as Delta-24-RGD.^4–6^ In patients with gliomas, MSCs can be injected intra-arterially via the carotid artery or its distal branches (middle cerebral and anterior cerebral arteries) or via the branches of the basilar artery or its branches (posterior cerebral artery) to enhance distribution and improve tumor targeting in diffuse or multifocal GBM.^7^ This strategy has the potential to improve therapeutic outcomes when surgical resection is either not feasible or is incomplete. Reliable non-invasive imaging of oncolytic virus-loaded MSCs is vital to assess their delivery efficacy, therapeutic activity, and safety in cell-based cancer therapies.^8,9^

Unlike invasive biopsies, non-invasive imaging could provide insights into *in vivo* cell biodistribution and persistence, track and quantify oncolytic MSCs, and enhance our understanding of their therapeutic effectiveness.^10^ Imaging techniques such as single-photon emission tomography (SPECT)^11,12^, positron emission tomography (PET)^13,14^, magnetic particle imaging (MPI)^15,16^, optical imaging^17,18^, and computed tomography (CT)^19,20^ have been investigated for cell-tracking applications. However, each approach presents distinct barriers to clinical translation, including potential injury to radiotracer-labeled MSCs and their therapeutic viral payload, limited tissue penetration with optical imaging, limited availability of human MPI scanners, and potential immune responses against genetically modified cells when reporter-based tracking strategies are used. In contrast, iron oxide particle (IOP) labeling enables MRI-based tracking of MSCs with high spatial resolution, broad anatomic coverage, and relatively low cellular toxicity, while avoiding the time constraints imposed by radionuclide decay.

Researchers have long investigated labeling mesenchymal stem cells with IOPs as a strategy for non-invasive *in vivo* cell tracking with MRI. IOPs generate strong MRI contrast detectable on standard clinical MRI systems, enabling repeated imaging without ionizing radiation.^21^ IOPs alter the longitudinal (R_1_) and transverse (R_2_/R_2_*) relaxation rates of surrounding water protons; however, they function predominantly as highly efficient transverse relaxation contrast agents. The magnitude of relaxation enhancement depends on particle size, magnetic properties, aggregation state, compartmentalization, and local concentration. These relaxation characteristics make IOPs particularly useful for cellular MRI, vascular imaging, and molecular imaging applications, where labeled cells or accumulated particles appear as regions of signal loss on R₂- and R₂*-weighted images.^21^ In preclinical studies, IOP-based MRI (IOP-MRI) has shown strong potential for cell-tracking applications. For example, Suryadevara et al. demonstrated that IOP labeling enabled MRI tracking of engrafted stem cells and assessment of therapeutic efficacy in cartilage repair.^22^ In another study, Wu et al. showed that IOP labeling enabled MRI-based tracking of chimeric antigen receptor (CAR) T-cell homing *in vivo* in a GBM model.^23^ Furthermore, in a recent study, MRI was used to track the migration of IOP-labeled MSCs toward the ischemic penumbra in a rat stroke model.^24^ Together, these studies support the use of IOP-MRI for longitudinal tracking of MSC biodistribution *in vivo*, particularly in oncology applications where MRI is already integrated into clinical care.

Achieving an optimal balance between MRI sensitivity, preservation of MSC function, and regulatory feasibility is critical for successful clinical translation of cell-tracking strategies. IOPs must be internalized in sufficient quantities to generate detectable MRI contrast, yet the labeling process must not compromise MSC viability, proliferative capacity, or multilineage differentiation potential. Over the past two decades, several strategies have been developed to enhance IOP uptake in MSCs while maintaining viability and function, including direct incubation, facilitated uptake using transfection agents to increase membrane permeability, nanoparticle complex formation to promote endocytic uptake, and physical membrane permeabilization techniques.^25–29^ In this study, we implemented a cell labeling protocol that combines facilitated uptake and nanocomplex formation to simultaneously load MSCs with IOPs and Delta-24-RGD oncolytic virus within a clinically feasible labeling time. We then assessed MSC integrity *in vitro* and quantified MRI detection sensitivity.

## Methods

In this study, we developed an integrated dual-loading protocol that enables concomitant loading of Delta-24-RGD and IOPs into MSCs by adapting established viral loading^30^ and IOP preparation^25,31^ strategies. To label MSCs, we used MegaPro IOPs (MegaPro Biomedical Co., Ltd., Zhubei City, Hsinchu County, Taiwan), a clinically translatable superparamagnetic iron oxide formulation that has successfully completed a Phase II clinical trial for MRI applications.^32^ MegaPro has been reported to exhibit higher transverse relaxivity (r₂) than several clinically approved or previously investigated iron oxide formulations, including ferumoxytol^22,27^, thereby providing greater per-cell MRI contrast sensitivity. Its high relaxivity enables detection of labeled MSCs at lower intracellular iron concentrations while maintaining a favorable safety profile, making it well suited for quantitative cell tracking studies.

### Cell Labeling process

Human bone marrow-derived mesenchymal stem cells (hBMSCs; Lonza, PT-2501) were labeled with the IOP MegaPro in the presence or absence of the oncolytic adenovirus Delta-24-RGD. Protamine sulfate (10 mg/mL; Fresenius Kabi USA, Lake Zurich, IL) was used as a cationic transfection agent to facilitate nanoparticle uptake, as previously described.^28^

Figure 1 illustrates the cell labeling scheme. On day 1, cells were seeded at 2.22 × 10⁴ cells/cm² in complete medium consisting of α-minimum essential medium (MEM, Cat. No. 15-012-CV, Corning, Glendale, AZ), 1% GlutaMAX^TM^ (Cat. No. 35050061, Gibco, Fisher Scientific, Waltham, MA), 5% human platelet lysate (PLTMax^®^, Cat. No. PLTMax100GMP, Mill Creek Life Sciences, Rochester, MN), and 2 U/mL heparin (10,000 USP units/mL, Fresenius Kabi). Post cell seeding, MSC cultures were incubated overnight at 37 °C in a humidified atmosphere containing 5% CO₂ to allow cell adherence. On day 2, 2 processes were initiated independently: 1) MegaPro medium preparation and 2) MSC virus loading. For MegaPro medium preparation, we prepared two equal-volume solutions in serum-free α-MEM: one containing protamine sulfate and another containing MegaPro Solution (see Supplementary Table 1 for volumes). We vortexed each solution, incubated it at 37 °C for 5 minutes, then combined them and incubated for an additional 5 minutes. The combined solution was transferred to an empty culture flask and incubated at 37 °C with 5% CO₂ for 4 hours on an orbital INFORS HT Celltron shaker system (25 mm orbit, 25 rpm, INFORS HT, Annapolis Junction, MD) to promote nanoparticle complex formation. In parallel, three hours after the start of the IOP complex formation step, the virus loading process was initiated; flasks containing adherent hBMSCs were gently washed once with pre-warmed magnesium and calcium free Dulbecco’s phosphate buffered saline (dPBS) at a volume equivalent to the culture medium volume to remove residual serum proteins that could impair labeling efficiency. The rinse was aspirated and replaced with serum-free medium equal to one-quarter of the total culture medium volume (e.g., 10 mL for a T225 flask with a final volume of 40 mL) containing Delta-24-RGD virus (multiplicity of infection (MOI) of 50), followed by agitation of the flask every 10 minutes during a 1 hour incubation period. Flasks with cells labeled only with IOP, i.e., without Delta-24-RGD, underwent the same process as above, except that the culture medium in the final 1 h step did not contain any virus. After the 1 h incubation step with or without virus, the previously prepared IOP complex medium was transferred to a 50 mL conical tube, and the remaining additives required to reconstitute the complete medium were added sequentially: heparin, GlutaMAX^TM^, and human platelet lysate, vortexing between each addition to avoid any precipitation. This medium accounted for three-quarters of the total culture volume when combined with the above additives (e.g., 30 mL for a T225 flask with a final volume of 40 mL). The volumes of protamine sulphate and MegaPro were selected (supplementary table 1) to achieve a final concentration of 10 μg/ml protamine sulphate and 100 μg/ml pf IOP in the complete medium. The complete medium, containing IOPs, was then added to the flask containing the cells, and all cultures were incubated for an additional 23 hours at 37 °C with 5% CO₂ on an orbital Celltron shaker, set to 25 rpm of continuous agitation to complete the single or dual cell labeling process. After the labeling time, the medium containing unincorporated virus and/or IOPs was aspirated, cells were washed with dPBS, dissociated using TrypLE^TM^ Express (Cat. no. 12604013, Thermo Fisher Scientific) and collected by centrifugation at 1,000 rpm. To remove any extracellular or non-internalized iron, cell pellets were resuspended in 10 mL of dPBS at room temperature, centrifuged, and the supernatant was aspirated for three washes. Cells were then resuspended at the desired concentration for downstream assays or experimental use.

**Figure 1:**
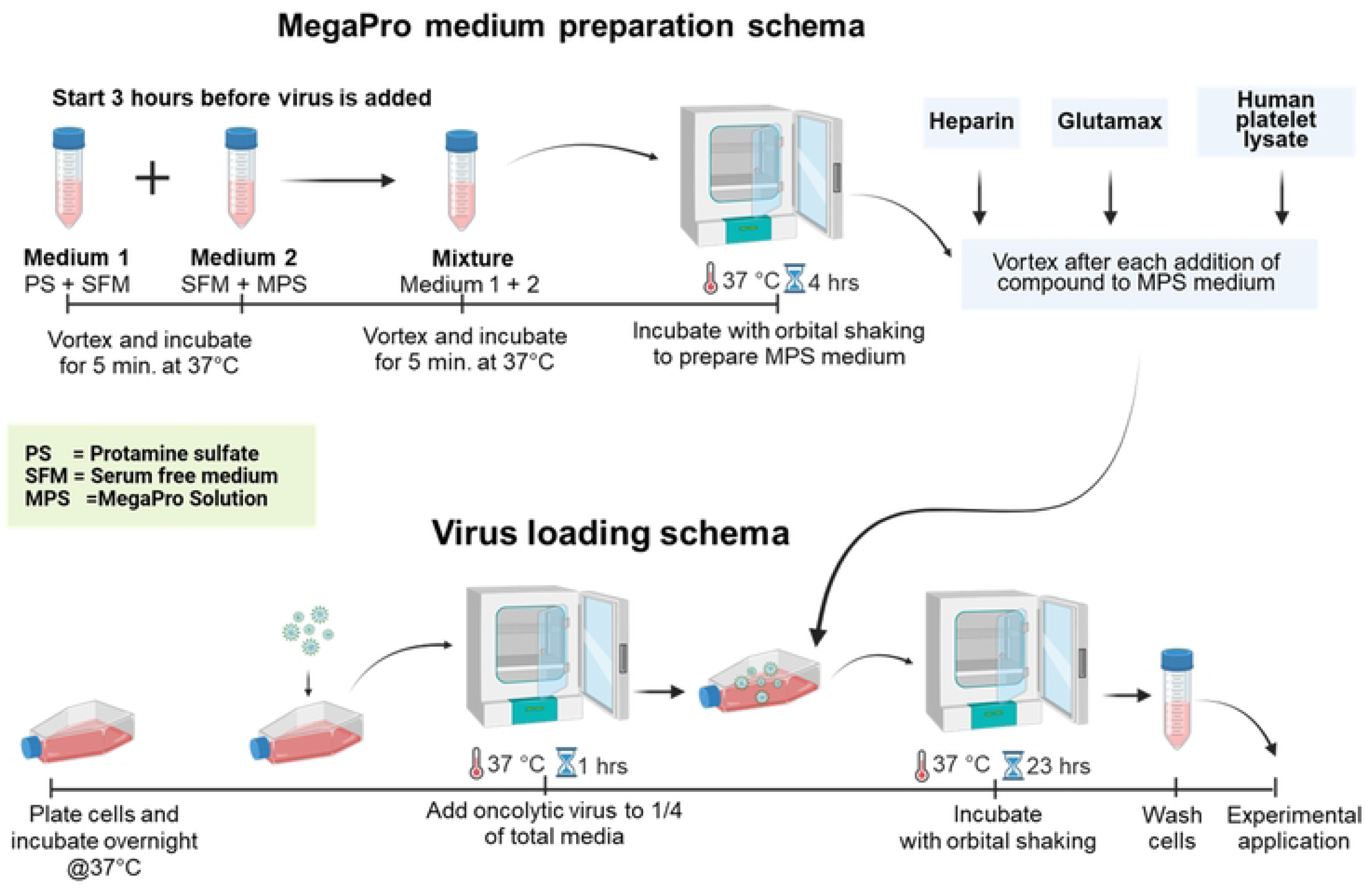
Schematic of the MSC labeling workflow for simultaneous dual labeling with the MegaPro iron oxide particles (IOPs) and virus, with respective steps for the IOP preparation (top) and the viral loading (bottom). After overnight MSC plating, Ad-Delta-24-RGD is added to the MSCs and incubated for 1 hour while in parallel, the IOP preparation begins 3 hours prior to the viral loading. Subsequently, the pre-prepared MegaPro IOP medium is added to the culture, and cells are incubated for an additional 23 hours to complete simultaneous viral and iron oxide labeling.

### Cell Viability

Cell viability was determined by Trypan Blue exclusion.^33^ Briefly, an equal volume of cell suspension was mixed with 4% Trypan Blue (Cat. No. T10282, Invitrogen, Thermo Fisher Scientific), and viability was quantified using a Countess 3 Automated Cell Counter (Cat. No. A49865, Invitrogen, Thermo Fisher Scientific) according to the manufacturer’s instructions. Cell viability was quantified for each condition in all assays at the completion of the labeling process (23 hrs after MegaPro medium is added to the cells).

### In Vitro Intracellular Iron Load Detection

Intracellular iron deposition was assessed by Prussian blue staining^34^. Briefly, Cells were seeded on poly-D-lysine (PDL)-coated coverslips (Cat. No. GG-15-PDL, Neuvitro Corporation, Vancouver, WA) and allowed to adhere prior to fixation. Cells were fixed for 20 minutes at room temperature in 4% paraformaldehyde (PFA; Cat. No. P6148, Sigma-Aldrich, Inc., St. Louis, MO), then washed with deionized (DI) water followed by dPBS. After washing, the coverslips were incubated for 30 minutes at room temperature with gentle rocking in a freshly prepared solution containing 5 mL of a 1:1 mixture of 10% Potassium hexacyanoferrate (II) trihydrate (Cat. No. P3289, Sigma-Aldrich, Inc.) and 10% hydrochloric acid (Cat. No. H1758, Sigma-Aldrich, Inc.), diluted in DI water.

Following staining, samples were washed thoroughly with DI water and counterstained with nuclear fast red (Cat. No. 24199C-250, Polysciences, Inc.,Warrington, PA) for 5 minutes at room temperature to visualize cell nuclei. After a final rinse with distilled water, coverslips were mounted using VectaMount AQ mounting medium (Cat. No. H-5501-60, Vector Laboratories, Inc., Newark, CA) and imaged using bright light microscopy. For each condition, a minimum of one coverslip was stained with Prussian blue during each assay to validate labeling.

### Colorimetric Iron Quantification Assay

We used a Prussian blue based colorimetric assay to quantify intracellular iron content in labeled cells.^35^ Control cells and cells obtained following the labeling process with and without the addition of Delta-24-RGD were resuspended in dPBS to a concentration of 1 × 10⁵ cells in a final volume of 50 µL. An equal volume of 6 N hydrochloric acid (HCl) was added to each sample, and samples were incubated in sealed tubes in a 37 °C water bath for 48 hours to facilitate iron extraction. Following incubation, tubes were centrifuged to pellet the cellular debris. The supernatant was transferred to white, glass bottom 96-well plates (Corning, Cat No.3903), and 50 µL of a 10% Potassium hexacyanoferrate (II) trihydrate solution diluted in DI water was added to each well. Plates were incubated for 15 minutes at room temperature, after which optical density was measured at 630 nm using a spectrophotometer plate reader (Agilent Technologies, BioTek Synergy H1). Iron content was quantified by comparison to a standard curve generated using known concentrations of iron prepared from the MegaPro contrast agent. All conditions were performed in technical triplicate and repeated across three independent biological experiments.

### Immunohistochemistry of Viral Load

Following iron labeling and viral transfection, cells were plated onto PDL coated coverslips and incubated overnight at 37 °C. Media was then aspirated, and cells were fixed in ice-cold methanol at −20 °C for 10 minutes. Coverslips were washed three times with dPBS containing 1% bovine serum albumin (BSA, Cat. No. A4503, Sigma-Aldrich, Inc.) and incubated in the final wash for 10 minutes for blocking. After aspiration, goat anti-Hexon primary antibody (Cat. No. AB1056, Sigma-Aldrich, Inc.), diluted 1:500 in dPBS + 1% BSA, was added and incubated at 37 °C for 1 h. Coverslips were washed three times with dPBS + 1% BSA and incubated for 30 minutes with a horse anti-goat polymer secondary antibody (Cat. No. MP-7405-50, Vector Laboratories, Inc.). After three dPBS washes, staining was developed for 10 minutes using DAB substrate (ImmPACT^®^ DAB Peroxidase (HRP) Substrate Kit, Cat. No. SK-4105, Vector Laboratories, Inc.), which reacts with horseradish peroxidase (HRP). We washed coverslips three times in dPBS and mounted them with VectaMount AQ mounting medium.

### Cell Invasion

We performed an invasion assay to determine whether the labeling process altered the migratory tropism and invasiveness of mesenchymal stem cells (MSCs) toward tumor-derived cues. We evaluated cell invasion through an extracellular matrix (ECM) barrier toward a chemoattractant using a transwell-based invasion assay, with methodological adaptations from a previously published protocol.^36^ Transwell inserts (6.5 mm diameter, 8 µm pore size; Cat. No. 3464, Corning) were coated with 100 µL of Matrigel (Cat. No. 356234, Corning) at a final concentration of 1.4 mg/mL and incubated for 1 hour at 37 °C and 5% CO₂ in a humidified incubator. MSCs were enzymatically dissociated using TrypLE^TM^ Express, resuspended in serum-free α-MEM, and 2 × 10^4^ cells were seeded into the upper chamber of each insert.

Three conditions were evaluated in labeled and unlabeled cells: (i) serum-free α-MEM in the lower chamber (negative control), (ii) α-MEM supplemented with fetal bovine serum (FBS; HyClone™ FetalClone™ I serum, U.S. origin, (Cytiva), Cat. No. 16777-232, VWR International, Radnor, PA) in the lower chamber (positive control), and (iii) U87-conditioned media in the lower chamber (experimental condition). U87-conditioned media were generated by culturing U87MG cells (Cat. No. HTB-14, ATCC, Manassas, VA) in serum-free media for 24 hours, then collecting and filtering the media through a 0.22 µm filter (Sartorius, 180E14) to remove cells and debris. Transwell plates were incubated at 37 °C and 5% CO₂ in a humidified incubator for 48 hours. Doubling time of the cells was observed to be approximately 48 to 72 hours in complete media; therefore, proliferation occurring during the experiment in serum-free media is expected to be negligible. All conditions were performed in technical triplicate and repeated across three independent biological experiments. We removed non-migrated cells on the upper surface of the membrane using a moistened cotton swab. Invaded cells on the lower surface were enzymatically dissociated and quantified using a Countess™ 3 Automated Cell Counter (Invitrogen). For qualitative, representative imaging, membranes from inserts selected to reflect typical invasion patterns were fixed with 4% paraformaldehyde for 30 minutes, rinsed with dPBS, and stained with crystal violet (Cat. No. V5265, Sigma-Aldrich,) for 10 minutes. Inserts were washed, air-dried for 24 hours, and imaged using light microscopy.

### Cell Differentiation

To assess the differentiation potential of iron-labeled stem cells compared to non iron labeled controls, we performed three lineage-specific differentiation assays: adipogenesis, chondrogenesis, and osteogenesis. We seeded cells in 24-well plates at densities of 1.9 × 10⁴, 9.5 × 10³, and 9.0 × 10⁴ cells per well, respectively. We replaced the differentiation medium every 3 days. We imaged differentiated cells using a Keyence fluorescence microscope (Keyence Corp., Osaka, Japan).

**Adipogenic differentiation** was assessed with a 14-day differentiation period using the StemPro™ Adipogenesis Differentiation Kit (Cat. No. A1007001, Thermo Fisher Scientific) according to the manufacturer’s instructions. Following differentiation, cells were fixed in 4% paraformaldehyde for 30 minutes, washed with dPBS, and subjected to two 3 minutes washes in 60% ethanol. Cells were then incubated in Oil Red O solution (Cat. No. O139, Sigma-Aldrich) for 15 minutes. After staining, coverslips were washed once with 60% ethanol for 3 minutes, followed by three, 3 minutes washes with DI water. Cells were counterstained with hematoxylin (Cat. No. 22-050-113, Expredia), rinsed with DI water.

**Chondrogenic differentiation** was assessed with a 14-day differentiation period using the StemPro™ Chondrogenesis Differentiation Kit (Cat. No. A10071-01, Thermo Scientific) according to the manufacturer’s instructions. Following differentiation, cells were fixed in 4% formaldehyde for 30 minutes, embedded in Fisher Healthcare Tissue-Plus™ O.C.T. Compound (Cat. No. 23-730-571, Thermo Fisher Scientific), and cryosectioned on to Superfrost Plus microscopy slides (Cat. No. 1255015, Thermo Fisher Scientific,) at 10 µm thickness. Slides were washed in dPBS, and stained with 1% Alcian Blue solution (Cat. No. TMS-010-C, Sigma-Aldrich (EMD Millipore)) for 30 minutes. After staining, slides were washed under running water for 2 minutes and then rinsed with DI water.

**Osteogenic differentiation** was assessed by using the StemPro™ Osteogenesis Differentiation Kit (A10072-01, Thermo Fisher Scientific,) according to the manufacturer’s instructions, with a 21-day differentiation period. Following differentiation, cells were fixed in 4% formaldehyde for 30 minutes, washed with DI water, and stained with 2% Alizarin Red S solution (TMS-008-C, Sigma-Aldrich (EMD Millipore)) for 3 minutes.

### Flow Cytometry

For flow cytometric analysis, a single suspension of unlabeled and MegaPro-labeled MSCs was stained with the fluorophore-conjugated antibodies CD105-PE (Cat. No. 323206, Biolegend), CD73-APC (Cat. No. 344006, Biolegend), and CD90-FITC (Cat. No. 389804, Biolegend) and respective isotype controls Mouse IgG1-PE (Cat. No. 400114, Biolegend), Mouse IgG1-APC (Cat. No. 400119, Biolegend), and Mouse IgG1-FITC (Cat. No. 400107, Biolegend). The viability dye Sytox Blue (Cat. No. S34857, Thermo Fisher Scientific) was used to exclude dead cells. Following staining, samples were acquired using CytoFLEX LX Flow Cytometer (Beckman Coulter Life Sciences, Indianapolis, IN). A minimum of 50,000 events were collected per sample. Data were analyzed using FlowJoTM software (FlowJo LLC, Salem, OR (Becton Dickinson Biosciences, now Waters^TM^ Biosciences)).

### Cell Preparation for Relaxvity MRI

Cells used to make the phantom were fixed in 4% paraformaldehyde for 30 minutes and washed with dPBS. We immobilized the desired cell numbers in a layer of 2% ultra-low-melting-point agarose (Cat. No. A2576, Sigma-Aldrich). We used fixed cells to make the phantom. The cell-containing layer was sandwiched between 2 layers of 3% agarose (Cat. No. A6013, Sigma-Aldrich,) inside of a 5-mm high-throughput NMR tube (Cat. No. 669995751, SP Wilmad-LabGlass, Vineland, NJ), shortened to approximately 5 cm length. Tubes were placed in a cylindrical, custom-designed holder containing slots for 8 tubes (sFigure 1). The total volume of the cell layer was 100 µl, and it was ∼5 mm thick. Seven cell sample were prepared with 1.5 x 10^5^, 1.2 x 10^5^, 9 x 10^4^, 6.1 x 10^4^, 3 x 10^4^, 1.5 x 10^4^, and 9 x 10^3^ MSCs in the cell layer, respectively. An 8^th^ sample contained no cells in the central 3% ULMP agarose gel layer, serving as background control.

### MRI acquisition and analysis

The acquisition parameters for R_2_-, R_2_*-, and R_1_-weighted MRI, acquired on a 7T MR system (Bruker Biospin, Germany) are given in Table 1. Five 1 mm thick slices covered the entire cell layer. The R_2_-, R_2_*-, and R1 parametric maps were computed from the MR images as previously described ^37,38^ using MATLAB (V2018) scripts (MathWorks, Inc., Natick, MA). Relaxivity maps were generated, with relaxivity defined as the change in relaxation rate between vials with and without IOPs normalized by IOP concentration. ROI analysis was then performed on the central 1-mm-thick slice through the cell layer, and mean ROI relaxivity values were calculated by averaging all voxels within each ROI. A single slice contained approximately 20 µL of cell layer volume, which is a typical voxel volume used in human brain imaging with R_2_* mapping. The minimum detection limit for each relaxation parameter was defined as the lowest cell density for which the mean relaxation parameter measured in the labeled-cell phantom exceeded the background agarose signal by at least three standard deviations (mean + 3 × Standard deviation of agarose estimates), based on measurements from the central imaging slice. For the cell MRI, ROI analysis results are reported as parameter estimates ± standard deviation of estimates.

**Table 1.** MR imaging parameters.

|  | TR (ms) | TE (ms) | echo spacing | FA<sup>(0)</sup> | No. of echoes | FOV (mm<sup>2</sup>) | Matrix size | Slice thickness (mm) | Slice gap (mm) |
| --- | --- | --- | --- | --- | --- | --- | --- | --- | --- |
| R<sub>2</sub><sup>*</sup> | 810 | 2.5 | 2.5 | 50 | 4 | 30x30 | 192x192 | 0.75 | 0.25 |
| R<sub>2</sub> | 2000 | 7 | 7 | 90,180 | 4 | 30x30 | 192x192 | 0.75 | 0.25 |
| R<sub>1</sub> | 1400, 1600, 2000, 2500, 3300, 4500 | 10 |  |  |  | 30x30 | 128x128 | 0.75 | 0.25 |

### Statistical analysis

We averaged technical replicates for *in vitro* biological experiments. Results are presented as the mean ± standard deviation from biological replicates. For estimated iron concentration from the calorimetric assay and cell viability analysis, we used Tukey’s multiple comparisons test. For the invasion analysis, we performed a paired Student’s t-test. We considered p < 0.05 significant. We performed statistical analyses in GraphPad Prism V11.02 (GraphPad Software, Boston, MA).

## Results

### In vitro MSC functionality is preserved after labeling

We investigated IOP loading *in vitro* in both IOP-labeled MSCs (MSC + IOP) and IOP + virus-labeled MSCs (MSC + IOP + virus) conditions. Compared with naïve unlabeled MSCs (Figure 2A and D), Prussian blue staining indicates consistent loading of IOPs across both IOP (Figure 2 B & E) and IOP + virus (Figure 2C & F) loaded MSCs. Further, Figure 2F shows colocalization between the oncolytic virus, highlighted by Hexon (brown), and the IOP visualized by Prussian blue (blue). Quantitative analysis revealed significant differences in intracellular iron between labeled and unlabeled conditions. Tukey’s multiple comparisons showed that average intracellular iron content was significantly higher in MSC + IOP (9.2 ± 2.6 pg/cell, p=0.0015) and MSC + IOP + virus (11.6 ± 1.4 pg/cell, p=0.0004) conditions than in naïve MSCs (0.1 ± 0.1). No significant difference in intracellular iron (P=0.2821) was observed between the two IOP labeled conditions.

**Figure 2:**
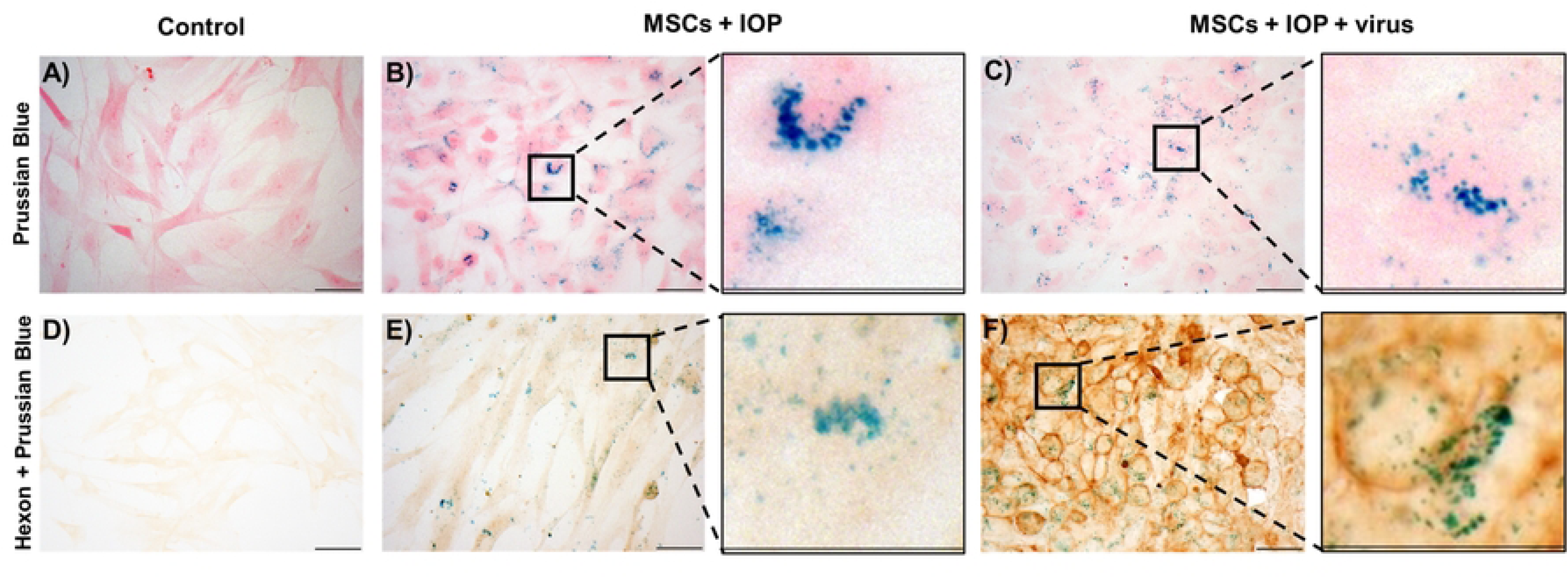
*In vitro* iron (Prussian Blue) and adenovirus capsid (Hexon, brown) staining. Panels A-C show Prussian Blue staining, while panels D-F show combined Prussian Blue and Hexon staining. A and D represent control conditions, B and E show MSCs + IOP, and C and F show MSCs + IOP + virus. Prussian Blue staining confirms iron uptake in both MSCs + IOP (B) and MSCs + IOP + virus (C) conditions, with no staining observed in controls (A). Dual Hexon and Prussian Blue staining demonstrates colocalization in MSCs + IOP + virus (F), with only background staining (no viral capsid) observed in control (D) or MSCs + IOP (E) conditions. Insets show zoomed-in regions of the indicated squares. Scale bar = 50 µm. White balance was applied to all images, and contrast was adjusted uniformly across images of the same staining using Fiji tool in ImageJ software.^49^

Cell viability remained high across all conditions, with no apparent reduction following labeling. Viability was 92.3 ± 5.5% in naïve MSCs, 93.0 ± 3% in MSC + IOP, and 91.0 ± 4.4% in MSC + IOP + virus conditions. In addition, MSCs retained their ability to differentiate into various cell lineages (Figure 3), including osteogenic, chondrogenic, and adipogenic lineages. Flow cytometric analysis confirmed that IOP-labeled MSCs retained high expression of the canonical MSC surface markers CD73, CD90, and CD105 (99.9% positive), comparable to unlabeled controls (99.9% positive), indicating that MegaPro labeling did not alter the immunophenotypic identity of MSCs (Figure 4).

**Figure 3:**
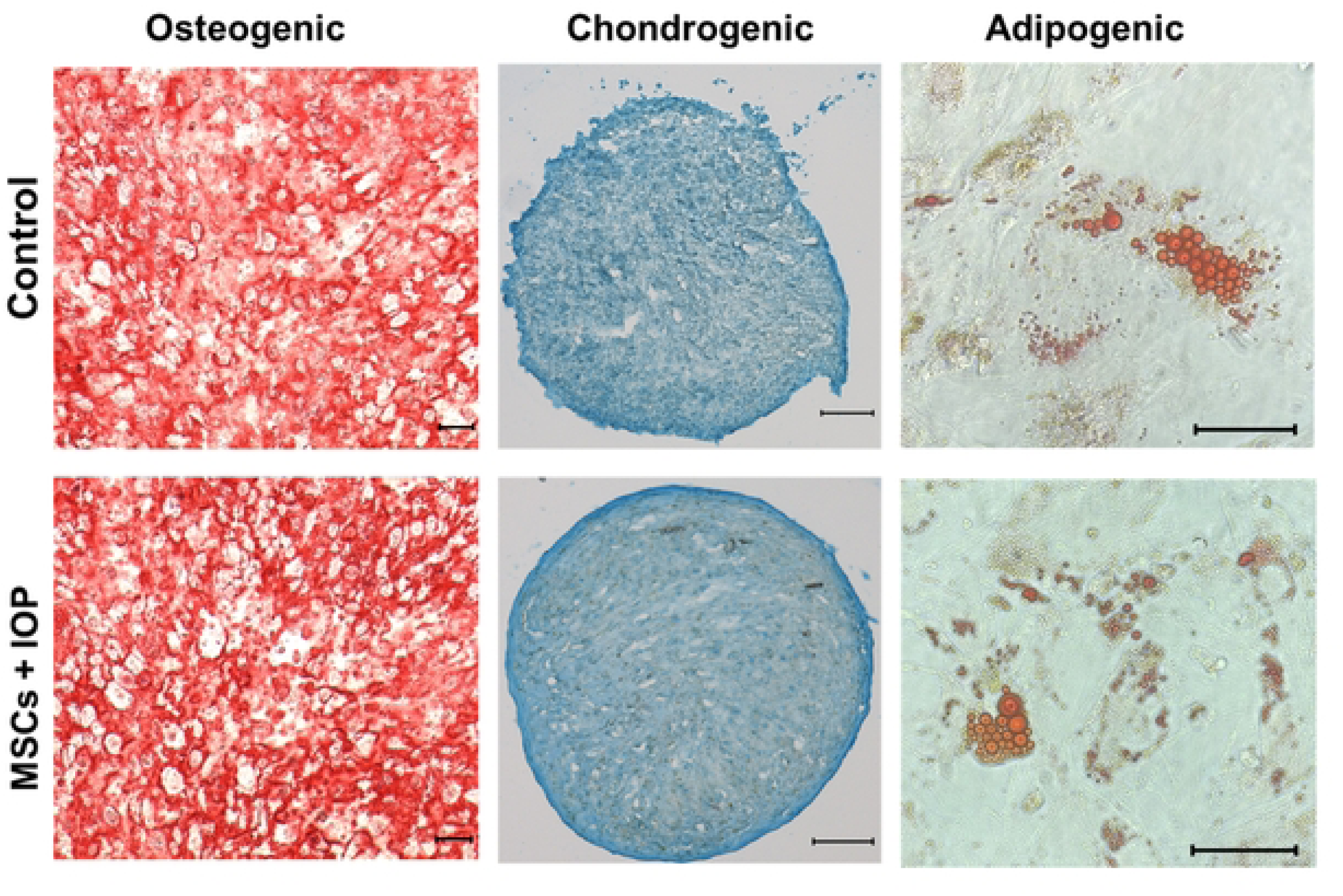
Osteogenic (left column), chondrogenic (middle column), and adipogenic (right column) differentiation of IOP-labeled MSCs. Similar patterns and ability to differentiate into the 3 cell lineages were observed for both, control (A) and MSCs + IOP (B). Scale bar 100 µm.

**Figure 4:**
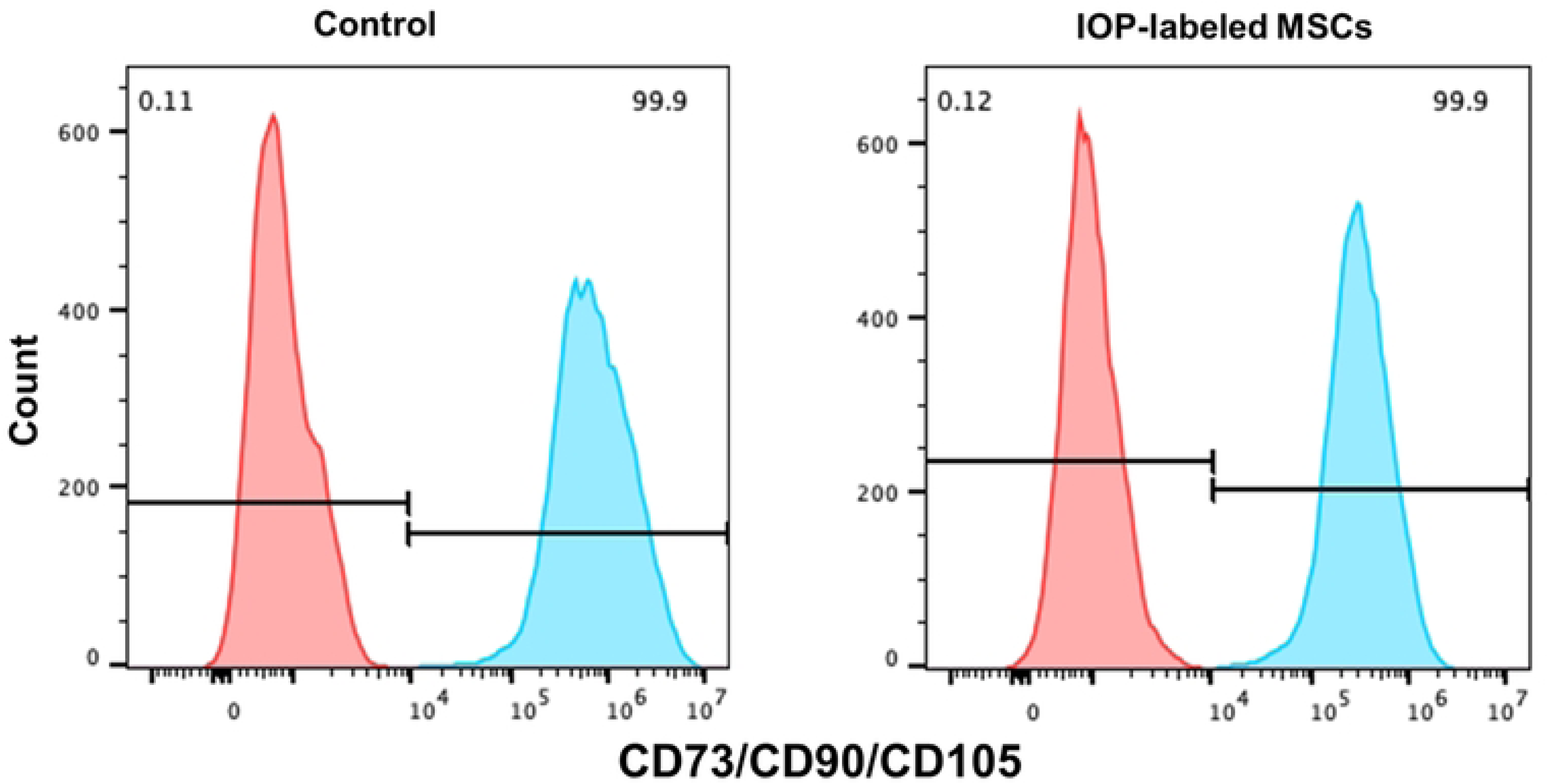
Flow cytometric analysis of A) Unlabeled human bone marrow-derived mesenchymal stem cells (BM-MSCs) and B) IOP-labeled MSCs shows that the expression of the canonical markers for MSCs is similar. Red peak (low intensity): Isotype control or unstained cells, defines background/autofluorescence baseline. Cyan peak (high intensity): MSCs positive for CD73/CD90/CD105. 99.9% gated positive confirms near-complete MSC marker expression in the unlabeled control, and IOP-labeled MSCs indicate that the cell population is a pure, well-characterized MSC preparation.

To evaluate the impact of IOP labeling on MSC migratory behavior, invasion assays were performed across three independent biological replicates. Across three independent biological replicates, MSC + IOP cells consistently exhibited reduced invasion compared with naïve cells both qualitatively (Figure 5A & B) and quantitatively (Figure 5C). Mean invasion values were 156,256 ± 35824 cells/50μl and 170,639 ± 18917 cells/50μl for MSC + IOP and naïve cells, respectively, corresponding to a modest 0.92-fold change. A paired t-test revealed that this reduction was statistically significant (p = 0.0082).

**Figure 5:**
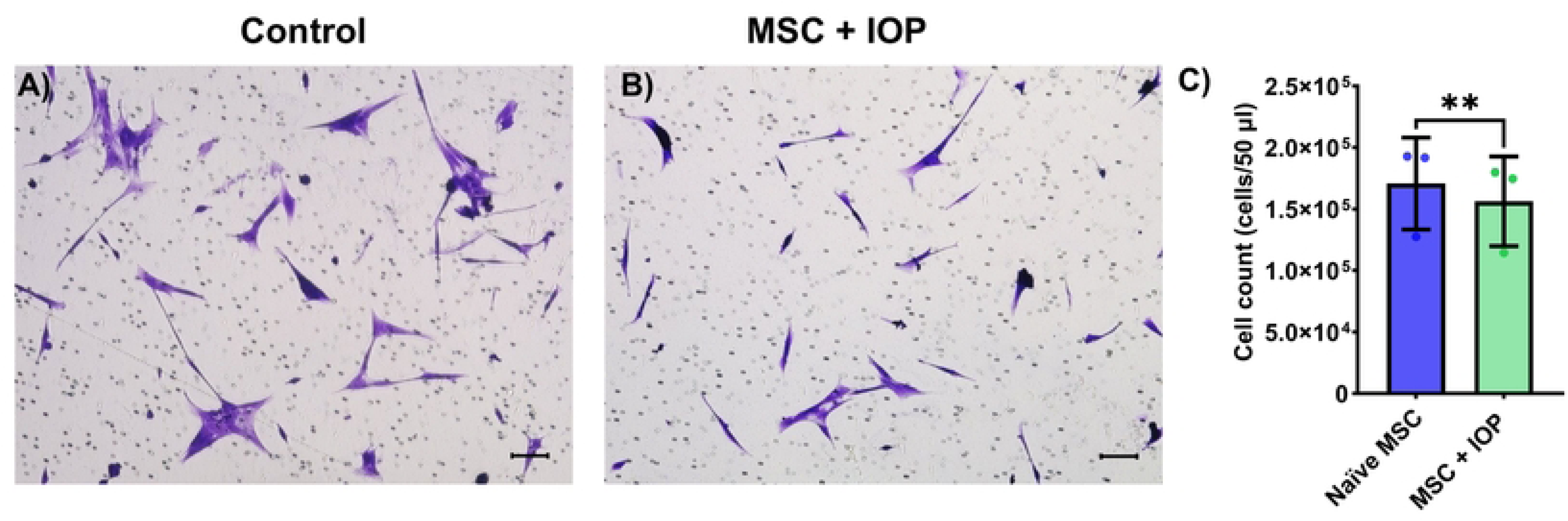
Qualitative and quantitative results for the invasion assay. A) Control (naïve MSCs) and B) MSCs + IOP-labeled MSCs with U87 medium chemoattractant. MSC+IOP cells exhibited significantly reduced invasion compared with naïve cells across three independent biological replicates (paired t-test, p = 0.0082). Mean invasion was reduced by 8.4% (0.92-fold change) both qualitatively (A and B) and quantitatively (c). Mean ± standard deviation of cell counts is shown in the bar plot. The scale bar is 100 µm in the qualitative images.

### Phantom studies

Phantom studies were conducted to further investigate the MRI sensitivity of IOP-labeled MSC cells across a range of cell densities. The generated quantitative parametric relaxivity maps (Figure 6A) and accompanying plots (Figure 6B) illustrate ROI analysis of the central 1-mm slice. For a phantom containing the highest cell number of 3×10^4^ cells, the relaxivities R_2_* (127.6 ± 42.0 s^−1^), R_2_ (77.1 ± 8.2 s^−1^), and R_1_ (0.55 ± 0.008 s^−1^) were found to be approximately 4, 2.5, and 1.5 times greater than the corresponding relaxivities for agarose (Table 2). Notably, the relaxation rate R_2_* declined when cell numbers reached or exceeded 18×10^3^ (131 ± 36.2 s^−1^), compared with lower numbers of 6×10^3^ (224.9± 11.5 s^−1^) and 12×10^3^ (229.8 ± 29.0 s^−1^) cells; however, it remained higher than values observed at both the minimum cell-density detection limits for R_2_* and agarose gel. The relaxation rates R_2_ and R_1_ showed a positive correlation with increasing cell numbers, reinforcing the relationship between cell concentration and magnetic resonance properties. The detection sensitivities for R_2_*, R_2_, and R_1_ in this experimental setup were approximately 2×10^3^, 6×10^3^, and 6×10^3^ cells per single slice (i.e ∼20 µl of the cell layer volume), respectively.

**Figure 6:**
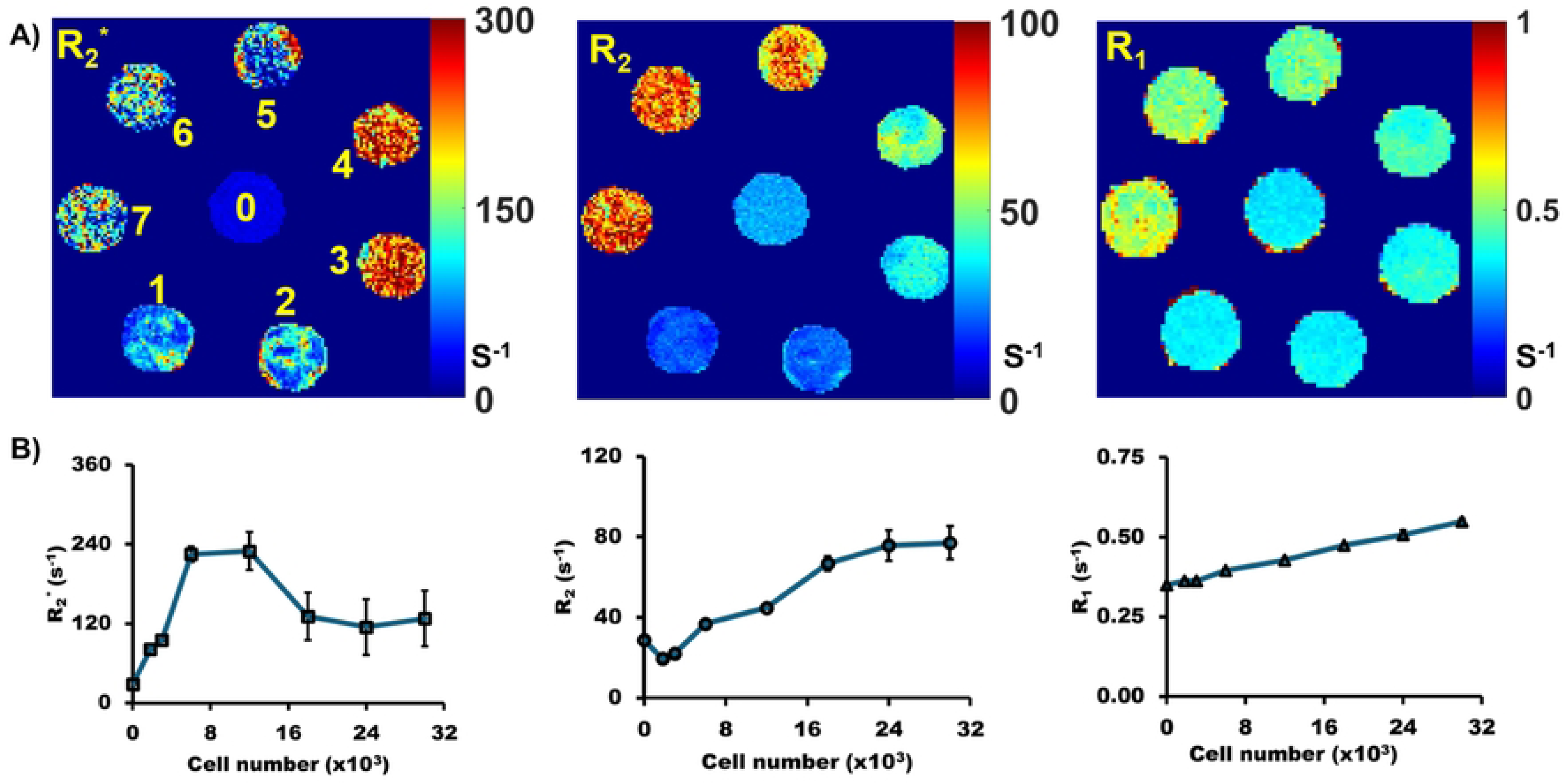
MR relaxation parameter results of IOP-labeled MSCs with different cell densities. A) Quantitative parametric maps of a 1 mm-thick central slice of the cell layer. B) Quantitative relaxation values measured from ROIs placed on the central 1-mm slice through the MSC layer, corresponding to approximately 20 µL of the total 100 µL phantom volume, across varying total cell numbers. Tube numbered 0 is the reference gel phantom (no cells). The cell sample tubes, numbered 1, 2, 3, 4, 5, 6, and 7, contain increasing cell numbers, i.e., 1.8 x 10^3^, 3 x 10^3^, 6 X 10^3^, 12 x 103, 18 x 103, 24 x 103, and 30 x 103 of IOP-labeled MSCs, respectively. Mean ± SD of the parameter estimates is displayed in the graphs.

**Table 2.** Quantitative (mean ± standard deviation) relaxivities based on ROI analysis for varying concentrations of IOP-labeled MSCs.

| Cell number ($\times 10^3$) | Agarose | 1.8 | 3 | 6 | 12 | 18 | 24 | 30 |
| --- | --- | --- | --- | --- | --- | --- | --- | --- |
| $R_2^* \text{ (s}^{-1}\text{)}$ | $28.7 \pm 0.4$ | $81.2 \pm 1.5$ | $95.1 \pm 1.1$ | $224.9 \pm 11.5$ | $229.8 \pm 29.0$ | $131 \pm 36.2$ | $114.9 \pm 41.9$ | $127.6 \pm 42.0$ |
| $R_2 \text{ (s}^{-1}\text{)}$ | $28.7 \pm 1.0$ | $19.5 \pm 0.9$ | $22.2 \pm 1.0$ | $36.9 \pm 1.5$ | $44.8 \pm 0.6$ | $66.8 \pm 3.5$ | $75.8 \pm 7.7$ | $77.1 \pm 8.2$ |
| $R_1 \text{ (s}^{-1}\text{)}$ | $0.35 \pm 0.001$ | $0.36 \pm 0.002$ | $0.36 \pm 0.005$ | $0.40 \pm 0.002$ | $0.43 \pm 0.002$ | $0.47 \pm .002$ | $0.51 \pm 0.014$ | $0.55 \pm 0.008$ |

## Discussion

For clinical translation of IOP-loaded MSCs as MRI-trackable carriers for Delta-24 oncolytic therapy, the labeling protocol must balance labeling efficiency with preservation of cell viability and function while remaining compatible with the viral loading timeframe used clinically. Therefore, in this study, we developed a dual-loading protocol to load both the IOP and the virus within a clinically feasible time frame. *In vitro* studies show excellent post-labeling viability, IOP loading into cells, and preserved functionality in IOP labeled cells. Phantom studies indicate that MRI can detect as few as 1,800 of the IOP-labeled MSCs in a 20 μl-volume phantom slice. Together, these findings support the feasibility of dual-loaded MSCs as MRI-trackable carriers for Delta-24-RGD therapy and provide a basis for future studies relating MRI signal to MSC biodistribution, cell dose, and therapeutic viral payload delivery *in vivo*.

Similar IOP loading was observed in both IOP- and IOP+virus-labeled MSCs, as demonstrated by qualitative Prussian blue staining (Figure 2B & C). Quantitative colorimetric analysis further confirmed comparable intracellular iron loading between groups, with virus-loaded MSCs exhibiting a slightly higher iron content (11.6 ± 1.4 pg/cell) than non-virus-loaded MSCs (9.2 ± 2.6 pg/cell). However, this difference did not reach statistical significance. The observed trend toward increased iron uptake in virus-loaded MSCs may reflect virus-induced modulation of cellular endocytic pathways, which have previously been implicated in nanoparticle internalization and intracellular trafficking.^39,40^ The iron loading achieved in our study exceeded previously reported values obtained using alternative methodologies, including mechanoporation IOP loading (0.94 pg/cell)^22^, passive diffusion (3 pg/cell)^27^, and noncomplex formation methods (<2 pg/cell).^25,26^ Compared with prior IOP-labeling strategies, these results suggest that the dual-loading approach can achieve higher intracellular iron loading while supporting MRI contrast generation and detection sensitivity. Furthermore, the observed colocalization of IOPs (Figure 2F; blue) and the oncolytic virus (Figure 2F; brown) supports the feasibility of dual functional cell labeling for combined imaging and therapeutic applications.

*In vitro* assessment further revealed that the IOP-loaded MSCs maintained preserved functionality. The preserved differentiation capacity of labeled MSCs, as demonstrated here, is critical for *in vivo* applications. These results are consistent with previous studies showing that carefully optimized IOP exposure does not significantly compromise MSC viability, migration, or differentiation.^41,42^ A modest reduction in invasion ability was observed for IOP-loaded MSCs compared to naïve MSCs, both qualitatively and quantitatively (a 0.92-fold change). Functionality retention after IOP labeling advances the translational readiness for MSC-based dual-delivery systems in cancer. Furthermore, despite dual manipulation, cell viability remained above 90%, supporting the biocompatibility of this labeling strategy. Collectively, these results demonstrate that MegaPro IOP labeling, either alone or in combination with oncolytic virus loading, preserves the key biological characteristics of MSCs, including viability, differentiation potential, and immunophenotype, while producing only a modest reduction in invasive behavior.

The ability to track MSCs *in vivo* depends on the MRI contrast, which is directly influenced by the number of IOPs internalized per cell. In this study, efficient labeling yielded a detection sensitivity of approximately 2 × 10^3^ IOP-labeled MSCs in a 20 µL volume, corresponding to a typical voxel size used in human brain imaging with R_2_* mapping. Direct comparison with prior studies is challenging, due to differences in iron oxide formulations, labeling protocols, imaging parameters, and reporting metrics (e.g., absolute cell number versus cell concentration). For example, detection thresholds on the order of 10⁴ cells have been reported by Castaneda *et al.*^31^, while Ohki *et al.* reported sensitivities of approximately 0.25 × 10⁶ cells/mL.^43^ In contrast, de Schellenberger *et al.* demonstrated detection at concentrations as low as 10³ cells/mL under optimized conditions.^44^ Despite these methodological differences, the detection sensitivity observed in this study falls within the range of previously reported values for MRI tracking of IOP-labeled cells.

As previously shown, *in vitro* MRI studies demonstrated that relaxation rates (R_2_*, R_2_, and R_1_) increase with cell density, reflecting the influence of superparamagnetic particle-laden cells on both transverse and longitudinal relaxation. R_1_ increased with cell density, although the magnitude of this effect was modest in comparison to the increases in the transverse relaxation rates. In contrast, R_2_ increased with cell density but plateaued at higher concentrations, indicating a more limited dynamic range under these acquisition conditions. R_2_* showed high sensitivity at low-to-moderate cell densities; however, at higher iron concentrations, R_2_* became nonlinear and, in some cases, showed apparent reductions. This behavior likely reflects susceptibility-induced field gradients that cause rapid signal decay before the earliest sampled echo, resulting in signal voids and underestimation of R_2_* under finite-echo-time sampling conditions. In this setting, R_2_ and R_1_ measurements may provide complementary information over concentration ranges in which R_2_* becomes unreliable.^45^ The use of ultrashort echo time (UTE) imaging may mitigate these effects by capturing rapidly decaying signals before severe dephasing occurs.^45–47^

This study has several limitations related to IOP-based cell labeling and MRI detection. First, this study lacks in vitro functional assessment of MSC behavior after viral loading, including differentiation assays. Second, our MRI studies used fixed cells, which may not completely replicate physiological conditions. Fixation eliminates active cellular processes and can alter intracellular water dynamics and compartmentalization, potentially affecting relaxation behavior and leading to an overestimation of MRI sensitivity.^15,48^ Furthermore, the fixed cell conditions lack physiological noise and motion artifacts, resulting in artificially high signal-to-noise ratios; consequently, the detection thresholds reported here likely underestimate the iron loading required for reliable *in vivo* visualization. Third, even though our protocol achieved excellent iron loading under clinically relevant constraints, it may be insufficient for robust *in vivo* detection. Although higher iron content could be achieved by optimizing labeling conditions, this would require careful evaluation of dose-dependent cytotoxicity and *in vivo* validation, which is outside the scope of this study. Finally, the relationship between relaxation rates and absolute cell numbers could not be established due to the nonlinear trends in the relaxation parameters.

## Conclusion

The findings of this study demonstrate that dual-labeling strategies for MSCs could enable MRI-based cell tracking while preserving oncolytic virus delivery capacity. The compatibility of iron labeling and virus-loading processes suggests broad applicability across cell-based immunotherapy platforms. Supported by advances in MRI relaxometry and MSC engineering, this approach holds strong translational potential. Future work will focus on *in vivo* validation, optimization of labeling protocols, and progression toward clinical translation.

## Acknowledgements

This work was supported by the Cancer Prevention and Research Institute of Texas (RR220038) and the Broach Foundation Fund for Neurosurgery Research. C. Chad Quarles is a CPRIT Scholar in Cancer Research. Special thanks to the MD Anderson Small Animal Imaging Facility (SAIF) supported by the NIH/NCI UT MD Anderson Cancer Center Support Grant P30 CA 016672. The authors thank MegaPro Biomedical Co., Ltd. for providing the iron oxide particles used in this study.

**sFigure1:** Illustration of a 3-layered cell phantom. Our cell phantom consists of a central layer with labeled MSCs encapsulated in 2% ultra-low melting (ULMP) point agarose (brown-colored layer) sandwiched between 2 layers of 3% ULMP agarose in a shortened 5 mm NMR tube.

## Notes

### Competing Interest Statement

The authors have declared no competing interest.

